# DNA Methylation Dynamics Reveal Unique Plant Responses and Transcriptional Reprogramming to Combined Heat and Phosphate Deficiency Stress

**DOI:** 10.1101/2025.11.19.689328

**Authors:** Alberto Lozano-Enguita, Victoria Baca-González, Adrián Morillas-Montáez, Jesús Pascual, Luis Valledor, Juan Carlos del Pozo, Elena Caro

**Author notes:** Corresponding authors: Elena Caro; Juan C. del Pozo.

## Abstract

Plants adapt to environmental challenges through epigenetic mechanisms that modulate gene expression without altering DNA sequence. Among these, DNA methylation is central to balancing genome stability and transcriptional flexibility. We analyzed methylation dynamics in *Arabidopsis thaliana* under heat, phosphate deficiency, and their combination—conditions that frequently co-occur in nature—using whole-genome bisulfite sequencing, small RNA-seq and RNA-seq of shoots and roots in a setup closely mimicking field conditions. Stress-specific patterns emerged: heat and combined stress led to CHH hypomethylation in both shoot and root, while phosphate deficiency triggered hypermethylation in shoots but hypomethylation in roots. Importantly, the epigenetic response to combined stress was not a mere additive effect of individual stresses but displayed a distinct methylation signature. While both RdDM and CMT2 pathways contributed to heat-induced changes, CMT2 predominated under phosphate deficiency and combined stress, underscoring mechanistic specificity. Methylation changes concentrated in transposable elements (TEs) and intergenic regions, yet TE methylation shifts showed limited correlation with TE or adjacent gene expression, suggesting methylation does not act as a direct transcriptional switch. Instead, stress-induced methylation may influence chromatin accessibility at regulatory regions, particularly transcription factor binding sites; GATA motifs appeared as especially relevant in our analyses. A striking signature emerged in the nuclear mitochondrial DNA (NUMT) region, where hypomethylation under heat and combined stress correlated with upregulation of oxidative phosphorylation genes, critical for thermotolerance. Our findings highlight DNA methylation as an intricate regulatory layer integrating environmental signals into plant adaptive responses, offering a foundation for strategies to harness epigenetic plasticity for crop resilience under climate change.

## Introduction

Abiotic stresses such as extreme temperatures and nutrient deficiencies hinder plant growth and yield by disrupting key physiological processes. As climate change intensifies the frequency and severity of these stresses, their impact on global agriculture is becoming increasingly significant. Gaining insight into how plants respond to these challenges is essential for breeding resilient crop varieties and safeguarding food security within the current environmental uncertainty.

Phosphorus is an essential macronutrient for plant growth and development. Although natural soils often contain enough phosphorus to withstand plant growth, much of it remains unavailable for uptake. To sustain high crop yields, modern agriculture is heavily dependent on the use of phosphate-rich fertilizers, a practice that is both economically unsustainable and environmentally damaging [1]. High temperatures are another environmental stressor that strongly affects plant growth, survival and yield [2]. Currently, climate change is leading to an increase in the frequency of extreme heat events, such as heatwaves, which can cause crop failure, reduced yields, and lower quality produce [3]. Heat waves rise temperature over the plant-optimal, increasing evapotranspiration and decreasing soil moisture. This limits nutrient availability, including inorganic phosphate (Pi), and restricts root uptake, ultimately causing complex and detrimental effects on plant–nutrient interactions [4]. In the context of climate change, crops are increasingly likely to encounter the combined stresses of high temperature and phosphorus deficiency, underscoring the importance of investigating plant responses to these combined stresses. Plant responses to combined stress are often not simply additive or linear combinations of responses to individual stresses. Instead, these responses are unique and complex, involving specific signaling pathways and regulatory networks, and should be considered for further studies on crop production and food security [5].

Epigenetics play an important role in plants response to environmental stresses by regulating gene expression without altering the underlying DNA sequence. This allows plants to rapidly adapt to changing conditions without genetic mutations [6]. Among epigenetic mechanisms, DNA methylation is particularly complex, as the same mechanism that enables the plasticity of DNA methylation for adaptive responses is also responsible for maintaining genome stability and ensuring consistent TE repression [7]. CG methylation is reliably inherited during DNA replication through the coordinated actions of DNA METHYLTRANSFERASE 1 (MET1) and the methyl-DNA-binding proteins VARIANT IN METHYLATION 1-3 (VIM1, VIM2, VIM3) [8]. Non-CG methylation in pericentromeric heterochromatin is maintained by three H3K9 methyltransferases (SUVH4, SUVH5, SUVH6) and CHROMOMETHYLASE 3 (CMT3) for CHG methylation, linking histone and DNA methylation [9]. CHH methylation is primarily maintained by CHROMOMETHYLASE 2 (CMT2) in heterochromatic regions and by DOMAINS REARRANGED METHYLTRANSFERASE 2 (DRM2) through the RNA-directed DNA methylation (RdDM) pathway in euchromatin [10]. The RdDM pathway establishes DNA methylation in all sequence contexts and maintains non-CG methylation in chromosome arms. RdDM involves two plant-specific RNA polymerases, Pol-IV and Pol-V, which are recruited to silent chromatin and produce small interfering RNAs (siRNAs) and long non-coding RNAs, respectively, that guide DOMAINS REARRANGED METHYLTRANSFERASE 2 (DRM2), to transposons and repeats based on sequence similarity, ensuring their proper methylation patterns [11].

DNA methylation has been demonstrated to regulate plant nutritional adaptations [12],[13]. In *Arabidopsis thaliana*, Yong-Villalobos et al. 2015 found that DNA methylation was crucial for establishing physiological and morphological responses to Pi starvation, as Pi deprivation resulted in changes in DNA methylation within genic regions linked to some Pi-responsiveness genes. Conversely, Secco et al., 2015 reported that the effects of Pi starvation on DNA methylation were primarily concentrated on TEs. Regarding the heat response, evidence from *Arabidopsis* presents a dynamic scenario. Initial acute heat exposure induces hypermethylation in certain chromosomal regions, followed by hypomethylation during recovery, with differentially methylated genes mainly associated with stress responses [14]. Early studies suggested the involvement of RdDM in the acute heat response, as mutations in *AGO4* or *NUCLEAR RNA POLYMERASE D2 (NRPD2*), a shared subunit of RNA polymerase complexes IV and V, led to increased heat sensitivity [15]. Subsequent observations highlighted CMT2-dependent CHH methylation as a significant factor in adapting to high temperatures. On one hand, CMT2-deficient *Arabidopsis* exhibited greater tolerance to both acute short-term and prolonged severe high and genome-wide association mapping connected *CMT2* with ecological temperature seasonality [16]. On the other hand, Jiang et al. (2024) revealed that CMT2 possesses a long, intrinsically disordered N-terminal region that compromises its protein stability, making it highly sensitive to elevated temperatures. Under heat stress, this structural feature leads to rapid degradation of CMT2, suggesting that plants may dynamically downregulate CHH methylation during thermal stress to allow transcriptional flexibility and activation of stress-responsive loci [17].

Despite the wealth of descriptive data on dynamic DNA methylation patterns in response to environmental stimuli, the underlying mechanisms remain poorly understood. It is crucial to comprehend these mechanisms in conditions as close as possible to real-world scenarios to develop crops with enhanced resilience to abiotic stresses, a goal that is becoming increasingly urgent in the context of global climate change.

## Results and Discussion

### Unique DNA methylation response to heat, phosphate deficiency and combined stress

Epigenetic responses to different abiotic stresses are highly specific with different stresses often producing contrasting and complex outcomes. For example, Pi deficiency increases global DNA methylation [18], whereas high temperature decreases it [14]. Interestingly, heat and Pi starvation response pathways are interconnected, as high temperatures modulate Pi starvation responses and expression of related genes [19]–[21]. To ensure our analyses closely reflect field conditions and enhance the applicability of laboratory results, we grew plants using the D-root system to keep roots in darkness (dark-grown roots), as light can trigger stress responses [22]. Additionally, for heat stress, plants were grown at 32 °C, a moderate heat, for an extended period to mimic real heat-wave conditions, and in the TGRooZ device, to form a temperature gradient in the root growing zone, mimicking natural soil conditions [21].

Plants exposed to elevated temperatures displayed fully functional roots, able to maintain a largely normal shoot morphology, even with increased fresh weight [21]. In contrast, under Pi starvation, dark-root plants exhibited a small decrease in root development and severe shoots phenotypes [23]. Under combined stress, plants formed a well-developed root system, but displayed shoot growth defects similar to those found in Pi starved plants (Figure 1AB).

**Figure 1.**
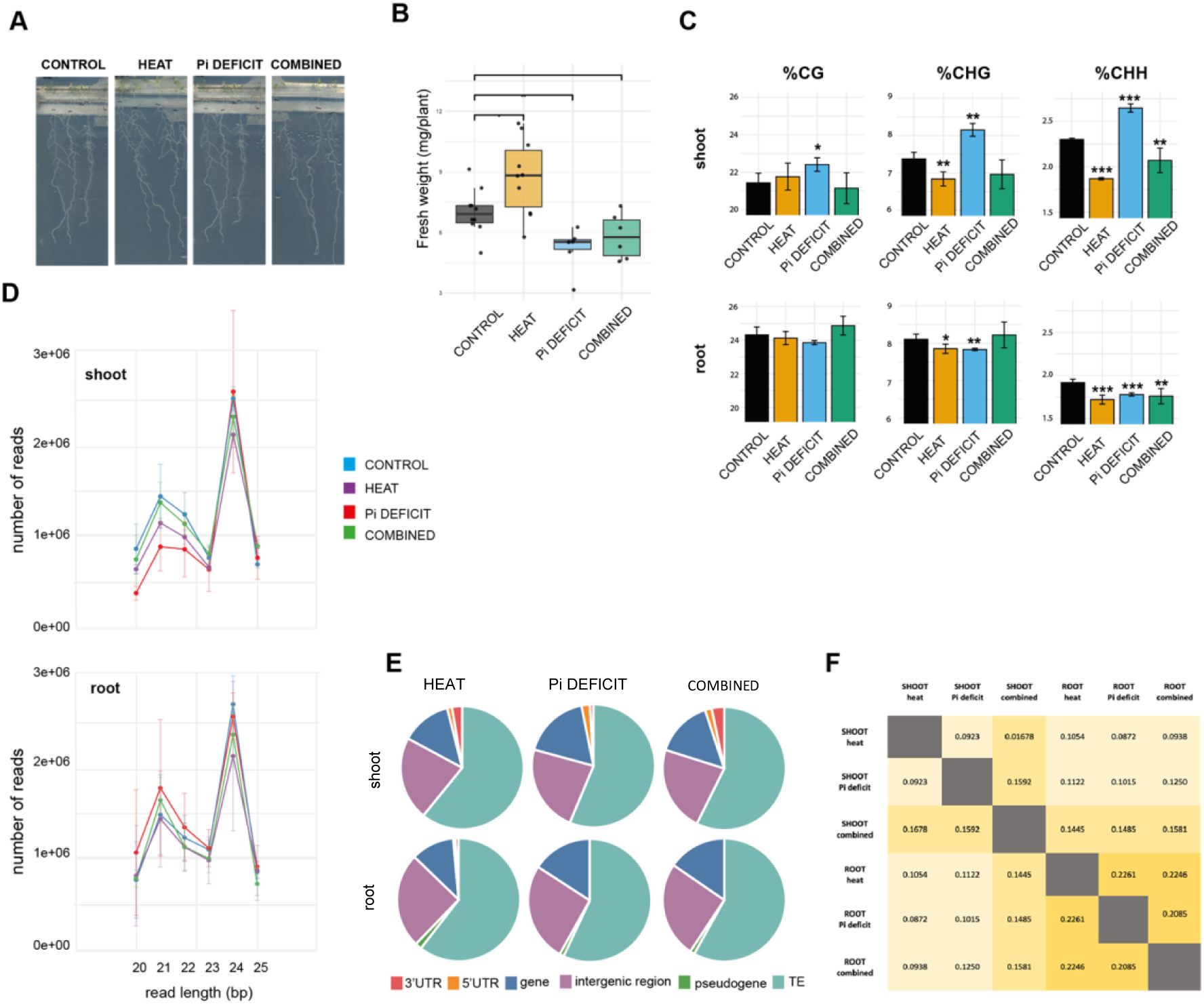
Effects of heat, Pi deficiency, and combined stresses on plant growth, DNA methylation, and small RNA profiles. **A)** Representative images of plants grown under control, heat stress, Pi deficiency, and combined stress conditions. **B)** Fresh weight per plant under the four treatments. Boxplots indicate median, interquartile range, and outliers. Asterisks denote statistically significant differences compared to control (*p < 0.05, **p < 0.01). **C)** Global DNA methylation levels in CG, CHG, and CHH contexts in shoots (top) and roots (bottom). Error bars represent standard deviation; asterisks indicate significance levels. **D)** Size distribution of small RNA reads (20–25 nt) in shoots (top) and roots (bottom). **E)** Genomic origin of small RNAs across treatments, represented as pie charts. **F)** Correlation matrix between DMRs in shoots and roots under different treatments.

Whole genome bisulfite sequencing allowed for analysis of the methylation level in each cytosine of the genome in both shoots and roots (Figure 1C). The results showed statistically relevant changes in non-CG contexts. Consistently with previously works, the shoots of plants subjected to heat showed a decrease in DNA methylation [14],[24] and in the case of Pi deficit, an increase in DNA methylation [18]. Interestingly, roots of plants grown under our unique setup conditions showed a global decrease in DNA methylation in all treatments. This result is contrary to previous works, where in response to Pi deficiency they reported an increase in DNA methylation [18]. This is likely because they grew seedlings with root illumination, a condition that generates an additive stress [22]. When plants suffered both stresses simultaneously, the response was not simply an additive combination of the responses to individual stresses, but instead it seemed to be a milder version of the response to heat. Both the shoots and the roots showed a decrease in DNA methylation, in this case specific only to the CHH context. Genome-wide profiling of DNA methylation levels (Supplementary Figure 1A) showed that the described general trends were true along all five Arabidopsis chromosomes, present in both their arms and centromeric regions.

To further explore the functional consequences of these methylation changes, we examined the dynamics of small RNA populations under the same stress conditions. Small RNAs, particularly 21-nt and 24-nt species, play key roles in gene regulation and epigenetic maintenance, making them an ideal proxy to assess whether stress-induced methylation shifts translate into altered silencing pathways. DNA hypomethylation in heat stress, Pi deficiency and combined stress in roots were all associated with an increase in 21-nt siRNAs, and no significant change in 24-nt siRNAs (Figure 1D). These trends suggest that stress-induced demethylation at regulatory regions promotes transcriptional activation, providing substrates for 21-nt siRNA biogenesis and enhancing post-transcriptional regulation of stress-responsive transcripts. In contrast, the abundance of 24-nt siRNAs, which are primarily involved in RdDM and heterochromatin maintenance, remained largely unchanged, consistent with the stability of transposable element silencing under these conditions. Interestingly, Pi deficiency in shoots showed the opposite pattern—hypermethylation accompanied by reduced 21-nt siRNAs—indicating transcriptional repression and diminished post-transcriptional gene silencing activity in this tissue. Together, these observations highlight a stress-and tissue-specific interplay between DNA methylation and small RNA pathways, where dynamic changes in 21-nt siRNAs suggest transcriptional changes.

In order to study these responses in more detail, we identified the differentially methylated regions (DMRs) for each tissue and treatment (Supplementary Figure 1B) and found that, in all cases, they were located mainly within TEs, with a high representation of intergenic regions and genes too (Figure 1E). We computed the Jaccard index between the DMRs in order to assess the extent of overlap between the elements that change methylation in the different tissues and treatments (Figure 1F). We found that the most similar response was between roots after the different treatments. In the case of the shoots, heat and combined stresses, which triggered a similar overall effect (decrease in DNA methylation), did not show high similarity, and surprisingly, the percentage of overlapping elements in DMRs was similar between combined and heat or combined and Pi deficit (Supplementary Figure 1C). Overall, our analyses underscore the specificity of the epigenetic response, with distinct patterns of hyper-and hypomethylation occurring in different genomic elements depending on tissue type and treatment.

### Molecular mechanism involved in the response

Interestingly, despite the specificity of the response, for all treatments and tissues the major changes regarding DNA methylation corresponded to the CHH context, a non-symmetrical methylation that must be re-established every cell generation by one of the de novo pathways: RdDM or CMT2 [8]. These two pathways are known to target different types of TEs, where RdDM primarily acts on small TEs in open, accessible and gene-rich regions, while CMT2 preferentially targets large TEs in constitutive heterochromatin [25]. Therefore, by analyzing the characteristics of the differentially methylated TEs under each stress condition and tissue, we could infer which pathway was predominantly affected, providing mechanistic insight into stress-induced epigenetic reprogramming. In the case of shoots exposed to heat, it seemed that the affected TEs were very similar in terms of superfamily distribution and length to those of the whole-genome TE population (Figure 2AB). For the remaining cases, however, the stress-induced DMRs were enriched in retrotransposons (Figure 2A) and long TEs (Figure 2B), suggesting a stronger effect on pericentromeric heterochromatin, where these elements are typically targeted for DNA methylation by CMT2 [26],[27].

**Figure 2.**
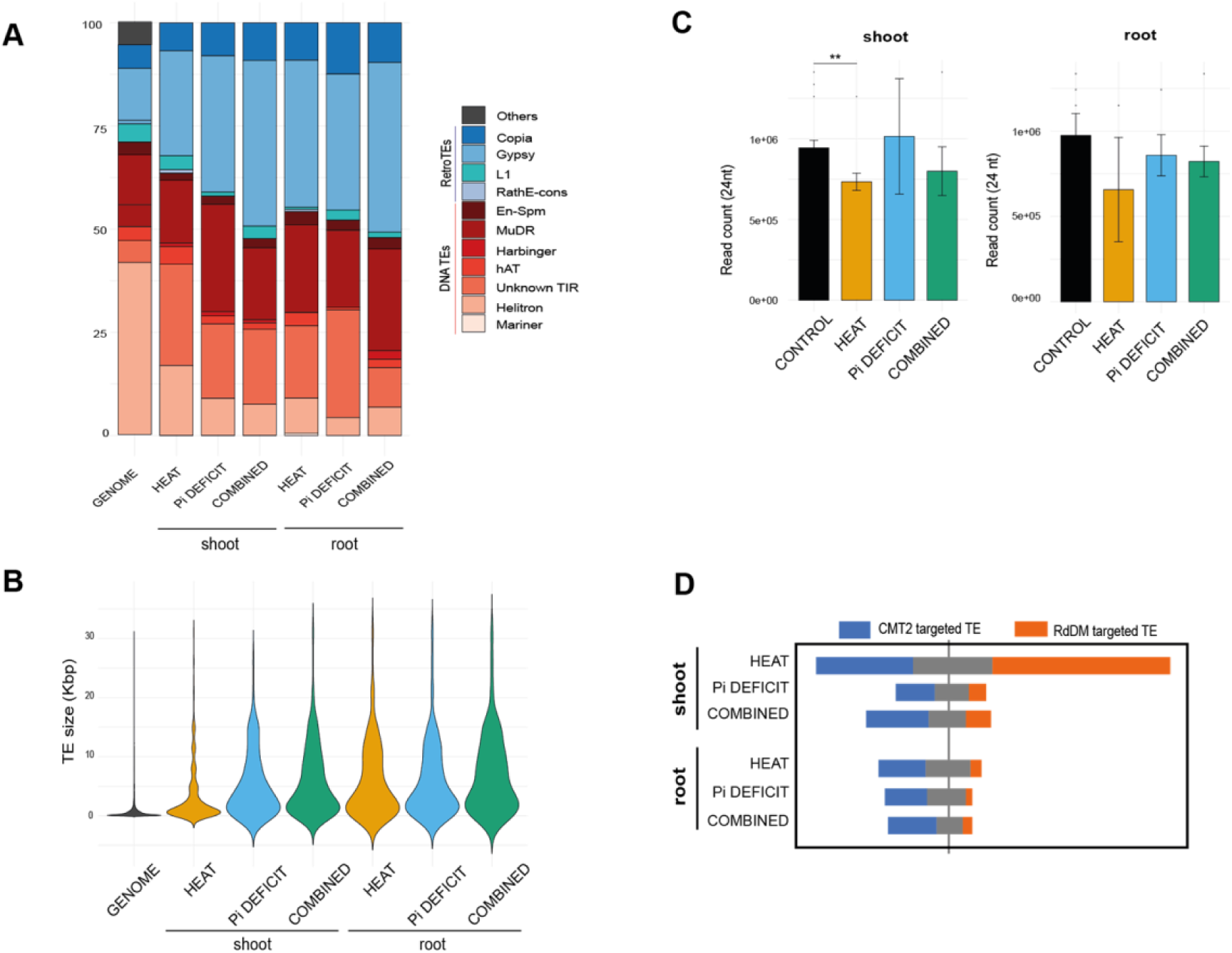
Transposable element (TE) composition, size distribution, and association with 24-nt siRNAs under heat, Pi deficiency, and combined stresses. **A)** Relative abundance of TE superfamilies in shoots and roots across treatments compared to the genome-wide distribution. Colors indicate TE classes, including DNA transposons (e.g., Helitron, Mariner) and retrotransposons (e.g., Copia, Gypsy). **B)** Violin plots of TE size distribution (kb) in shoots and roots under different treatments. **C)** Abundance of 24-nt siRNAs mapping to TEs in shoots (left) and roots (right). Error bars represent standard deviation; asterisks indicate significant differences compared to control (**p < 0.01). **D)** Proportion of TEs targeted by CMT2-mediated CHH methylation (blue) versus RdDM-mediated CHH methylation (orange) in shoots and roots under stress conditions.

Our TEs data indicate that different mechanisms are involved in these responses, likely involving changes in RdDM and/or CMT2 activity, or shifts in the balance between methylation and demethylation. To further explore the involvement of the RdDM pathway, we analyzed small RNA sequencing data generated from the same tissues and stress conditions focusing on 24 nt siRNAs. Consistently, the accumulation of siRNAs in TEs was only altered in shoots under heat stress, while it remained unaffected in other tissues and treatments (Figure 2C), supporting the conclusion that RdDM plays a role specifically in the shoot response to heat stress. In order to further validate these observations, we took advantage of previous studies where Arabidopsis TEs were characterized as CMT2-targeted or RdDM-targeted, depending on their demethylation in either *cmt2* or *drm2* mutants [28]. We analyzed the data from our DMRs according to this information (Figure 2D), and, consistently with our previous observations, the shoot response to heat involves TEs dependent on both CMT2 and RdDM. In contrast, the response to other stress conditions and in other tissues was clearly governed by CMT2. These findings reinforce the idea that epigenetic responses to abiotic stress are highly specific, not only in terms of the direction and magnitude of DNA methylation changes (hypo-or hypermethylation), but also regarding the molecular pathways responsible for establishing these changes. This highlights the differential behavior of plant organs and the unique nature of combined stress responses, where distinct mechanisms are selectively activated depending on the context.

Importantly, although previous studies reported rapid degradation of CMT2 protein at high temperatures (37 °C) [9] and transcriptional regulation of RdDM components under heat stress [29], we did not observe neither a decrease in CMT2 protein in TGRooZ seedlings (Supplementary Figure 2), nor clear transcriptional changes in key players of these pathways, apart from slight differences in ROS1 under Pi deficiency, likely due to the methylation-sensitive regulatory element in its promoter that controls ROS1 transcription, or in DML2 under combined stress in roots. Therefore, while our data strongly suggests that regions dependent on RdDM and CMT2 are involved in stress-induced methylation changes, the precise regulatory basis—whether through altered methylation, active demethylation, or both—remains unclear. These observations emphasize that the mechanisms underlying DNA methylation dynamics in response to abiotic stress may differ depending on growth conditions and stress combinations, and should be addressed in future studies.

### Effect of DNA methylation changes on TE activation

As mentioned before, TEs appeared to be the main targets of DNA methylation changes in response to stress (Figure 1E). Their overall methylation levels followed the same trends observed for global DNA methylation, with a marked increase in shoots under Pi deficiency, and a decrease in both shoots and roots under heat and combined stress conditions (Figure 3A). Because changes in TE methylation can influence their transcriptional activity, we analyzed RNA-seq data from plants grown under the same conditions used for DNA methylation profiling (Figure 3B). Although many TEs showed altered expression in response to stress, this was not generally correlated with changes in their DNA methylation status, as only a small subset of differentially expressed TEs were also differentially methylated. These observations are consistent with previous reports indicating that, after heat treatments, while DNA methylation contributes to TE silencing, it is not necessarily the primary or sufficient factor for their reactivation [30],[31] and that in the Pi starvation response, no clear relationships were found between TE differential expression and differentially methylated regions (DMRs) in either Arabidopsis or rice [13],[18].

**Figure 3.**
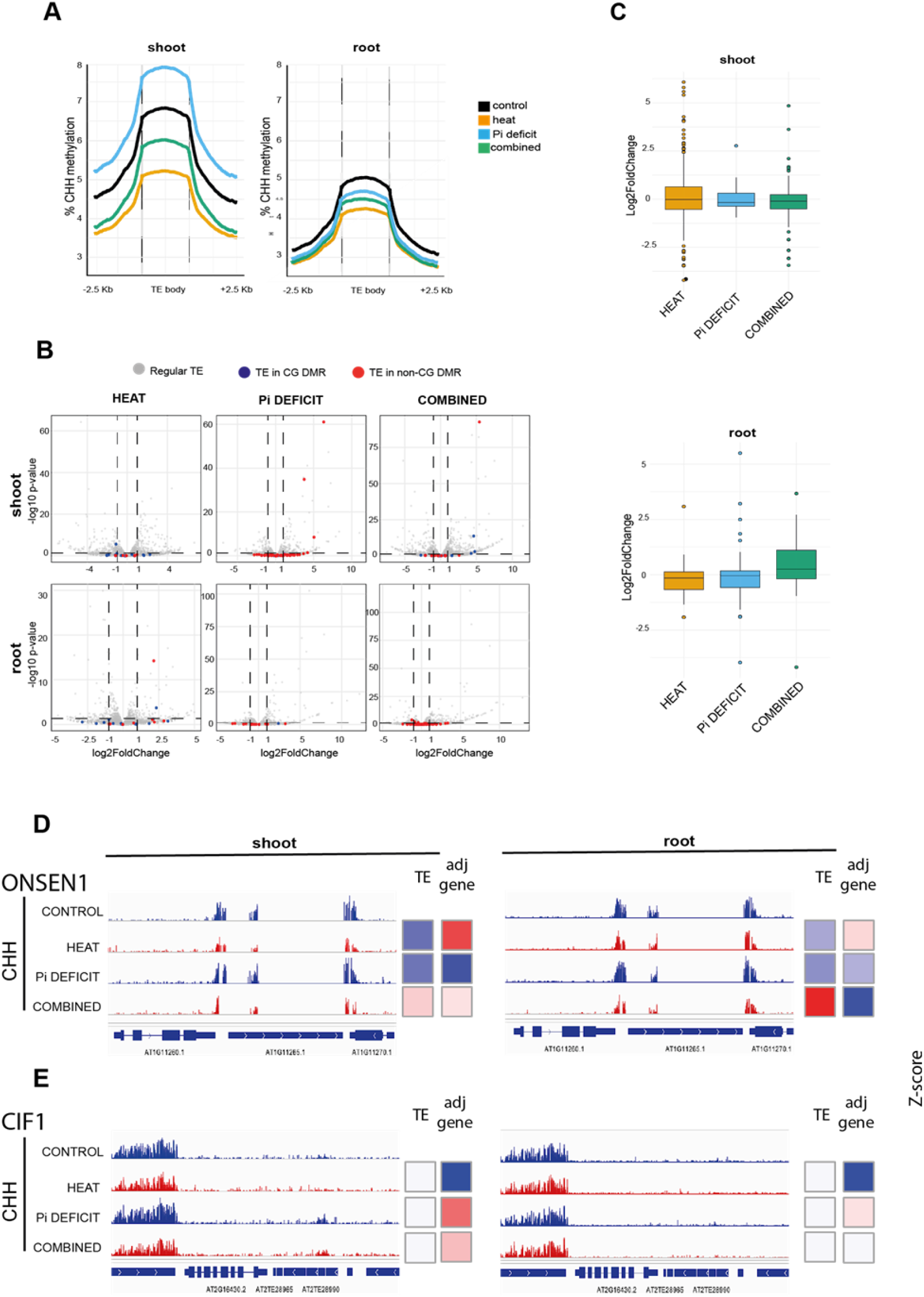
Relationship between DNA methylation and expression at stress-responsive transposable elements. **A)** Metagene plots showing DNA methylation levels across Tes and flanking regions in shoots (left) and roots (right) under control, heat, Pi deficiency, and combined stress conditions. **B)** Volcano plots of Tes in shoots (top) and roots (bottom) for heat, Pi deficiency, and combined stresses. Red dots indicate TEs in non-CG DMR; blue dots indicate Tes in CG DMR. **C)** Boxplots showing expression of genes adjacent to TEs in DMRs in shoots and roots. **D–E)** Genome browser views of representative loci: **ONSEN1** (D) and **CIF1** (E), showing DNA methylation tracks and expression of differentially methylated Tes and adjacent gene under different treatments.

### Effect of DNA methylation changes on gene expression

It is well established that changes in the epigenetic marks of TEs can contribute to stress adaptation, not only by affecting their own expression but also by influencing the expression of nearby genes through cis-regulatory mechanisms [32]. To test whether this occurs under our stress conditions, we analyzed the expression of genes located adjacent to differentially methylated regions (DMRs). We did not observe significant changes in expression across the overall set of neighboring genes (Figure 3C), suggesting that, in our experimental setup, stress-induced changes in TE DNA methylation are not directly associated with transcriptional changes—neither in the TEs themselves nor in their adjacent genes.

To complement the global analysis, we next examined representative transposable elements previously reported to be responsive to stress (Figure 3DE and Supplementary Figure 3AB). Regarding heat, these included members of the ONSEN family; ONSEN1 (Figure 3D) and ONSEN2 [33], the TE located in the promoter region of *CML41* (AT3TE76530) [29] and full-length Copia-35 elements such as *AT1TE51360* [34] (Supplementary Figure 3A). For these loci, we assessed both DNA methylation and transcriptional responses under our stress conditions and additionally evaluated the expression of adjacent genes to explore potential cis-regulatory effects. This targeted approach allowed us to confirm that heat and combined stress treatments lead to a decrease in CHH DNA methylation, but its correlation with the expression of TEs and neighboring genes is not direct. Previous works, where overexpression of these TEs and genes had been characterized, used much stronger heat treatments that were homogenously applied to the shoots and roots. Our milder experimental conditions were sufficient to trigger the DNA methylation changes previously reported; however, transcriptional alterations were evident only under combined stress. This raises the compelling possibility that stress severity plays a critical role in determining the extent of epigenetic modifications and also their functional translation into transcriptional outcomes.

Regarding Pi deficit, we analyzed the effect on gene expression of changes in DNA methylation in the TEs located upstream of CIF1 (Figure 3E), PAP10, AT5G22300 and AT4G01820 (Supplementary Figure 3B). These are all phosphate-responsive genes reported to show differential methylation and expression in response to Pi starvation [12],[18]. Here, it is interesting to highlight the differences between shoot (characterized by an increase in DNA methylation) and roots (characterized by a decrease in DNA methylation), a result reported for the first time in this study, probably due to the fact that in our setup, the additive stress generated by the root illumination was avoided by the use of the D-Root. In this case, there is a certain correlation where hypermethylation in TEs in the promoters of these genes led to their repression in shoot tissues. However, in roots of Pi deficient seedlings, TEs were not hypermethylated and adjacent genes seem to be induced (Supplementary Figure 3B). All these data suggest that the regulatory landscape is complex, and individual epigenetic changes—such as DNA demethylation—do not directly determine gene expression outcomes. Rather than acting as a direct ON/OFF switch, the loss of DNA methylation may function by allowing chromatin to become accessible and transcriptionally competent if additional regulatory signals are present.

Consistent with this idea, previous studies have suggested that changes in DNA methylation in intergenic regions could control the availability of TF binding sites to reprogram transcriptional networks under abiotic stress [35]. Methylation patterns are often associated with closed chromatin, what could prevent TF binding and activation of expression in these regions. To gauge this hypothesis, we analyzed the presence of TF binding sites within differentially methylated regions in intergenic regions after heat and Pi deficit. We found that, regardless of the stress and the tissue, DNA methylation changes after stress occurred preferentially over TF binding sites of certain families (Figure 4A). This was the case for Heat Shock Factor binding sites in heat (shoot and root) and combined (shoot), as expected due to the upregulation of HSP targets in these conditions and validating the analysis. Other known stress responsive TFs like TCPs [36],[37], SBPs [38] and ARR-B [39]–[41], also showed up as enriched in DMRs (Supplementary Figure 4). However, the strongest binding site overrepresentation was that of GATA TFs. GATA are a family of 30 genes in *Arabidopsis thaliana*, previously reported to play key roles in regulating developmental processes and environmental responses, particularly through their involvement in light signaling, nutrient sensing, and hormonal pathways [42],[43]. GATA transcription factors exhibit strong sensitivity to methylation in their binding site, with binding to their motifs markedly reduced when cytosines are methylated [44], indicating that DNA methylation changes in response to stress could be effectively blocking or allowing access GATA TF to regulatory sites and thereby, influencing transcriptional regulation. RNAseq data showed that GATA family members are mostly upregulated in shoots and roots of stressed plants (Figure 4B), supporting a possible contribution of GATA gene regulation in the response to heat and Pi deficit that should be further investigated in the future.

**Figure 4.**
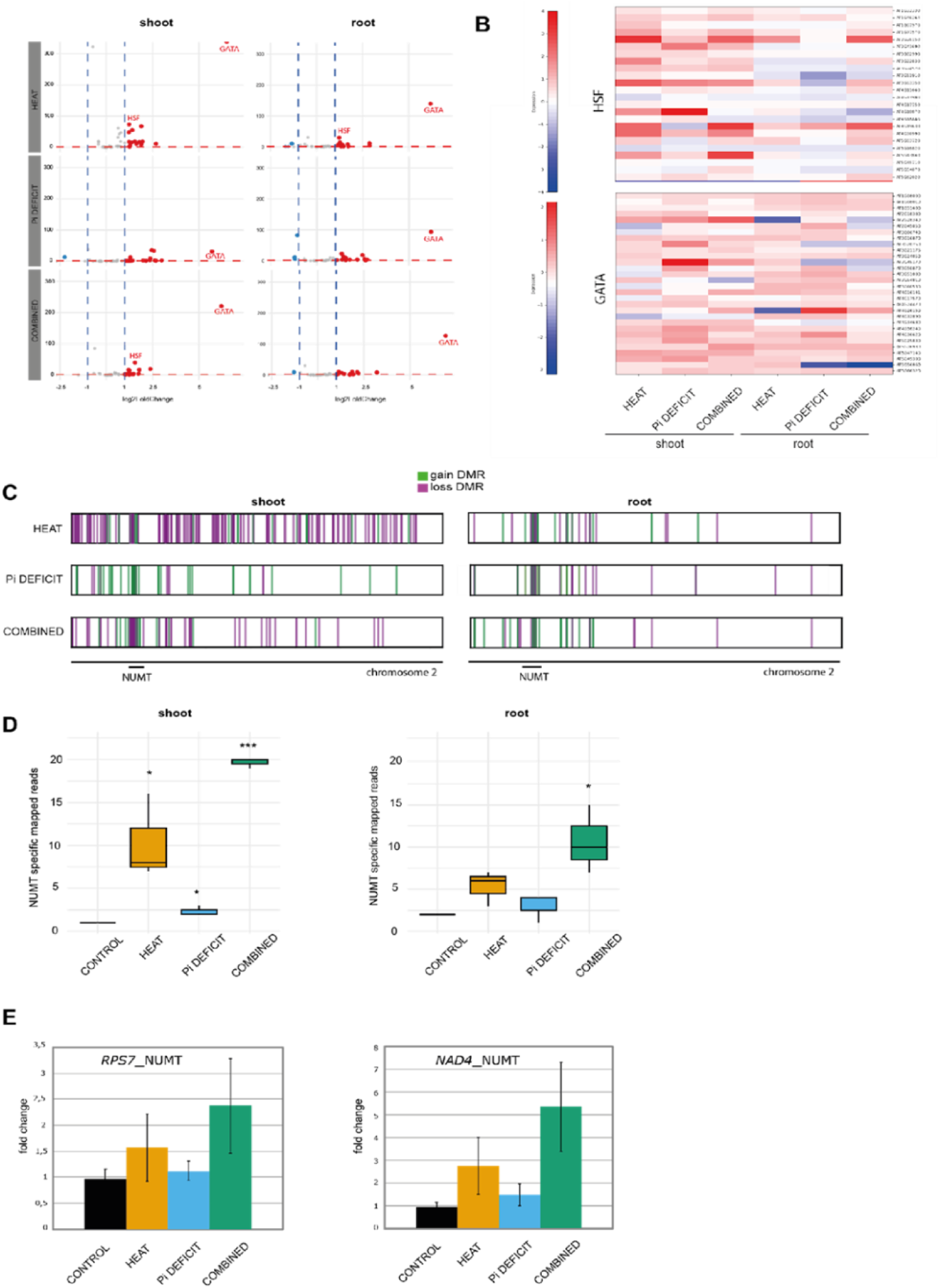
Relationship between DNA methylation and expression at stress-responsive intergenic regions and genes. **A)** Analysis of enrichment of transcription factor binding sites at intergenic DMRs under control, heat, Pi deficiency, and combined stress conditions. **B)** Heatmap showing expression of heat shock factors (up) and GATA transcription factors (down) under control, heat, Pi deficiency, and combined stress conditions. **C)** Distribution of DMRs in shoots and roots under different stress conditions along chromosome 2. Green marks hypomethylated regions; purple marks hypermethylated regions. **D)** Boxplots showing specific NUMT gene abundance in shoots and roots from RNAseq data. **E)** Bar plots showing specific NUMT gene abundance in shoots from qPCR data.

In a final effort to study the effect of changes in DNA methylation on gene expression, we focused our study in the DMRs located within genes. Interestingly, we noticed that most of the genes whose DNA methylation changed in response to stress localized in a very specific region in the short arm of chromosome 2, near its centromere. On closer observation, we could identify this region as the largest insertion of mitochondrial DNA in the Arabidopsis genome [45]. Nuclear mitochondrial DNA segments, known as NUMTs, are generally considered non-functional remnants of past organelle-to-nucleus DNA transfer events. A more detailed analysis of our data showed that chromosome 2 NUMT genes were hotspots of DNA methylation changes after any kind of stress and in both tissues, shoots and roots. In the case of heat and combined stresses it is clearly hypomethylated, and for Pi deficit, hypermethylated in the shoot, and hypomethylated in the root (Figure 4C).

To study the effect of these epigenetic changes on transcription, we used our RNAseq data. However, since NUMT shares 99.933% sequence identity with the mitochondrial genome, distinguishing reads originated from nuclear versus mitochondrial sequences becomes challenging, especially when centromeric regions in the Araport11 reference exhibit low sequencing quality, complicating downstream genomic analyses. In order to determine differences between NUMT and mitochondrial genome genes, we used recent long-read sequencing data [45] which allowed us to identify single-nucleotide polymorphisms and structural variants unique to each. We then measured the RNAseq reads that mapped specifically to NUMT and analyzed their accumulation (Figure 4D). Interestingly, for NUMT genes, there was a clear correlation between DNA methylation and expression. The decrease in DNA methylation (in heat and combined stresses) was concomitant with transcriptional upregulation. In the case of Pi deficiency in shoots, where NUMT region is hypermethylated, its genes remained silent (Figure 4D). These results were validated using specific qPCR primers able to discern between NUMT/mitochondrial genes in order to specifically measure the expression pattern of NUMT genes after stress. Our results for 2 example genes (Figure 4E) clearly show how the loss of DNA methylation in the chromosome 2 NUMT region correlates with a specific upregulation of gene expression. Reinforcing this data, we observed that *cmt2* plants, which have an overall lower level of CHH DNA methylation [27], show enhanced upregulation of NUMT genes upon heat stress [31].

Plant respiration and ATP synthesis via mitochondrial oxidative phosphorylation (mtOXPHOS) has been identified as the pathway with highest correlation to heat stress response in Arabidopsis [46]. Moreover, impaired respiration resulting from mtOXPHOS dysfunction is associated with increased thermosensitivity in plants, suggesting its importance for plant survival under heat stress. It is tempting to speculate that the observed increase expression in NUMT-encoded OXPHOS genes in response to heat could confer an adaptive advantage, potentially enhancing plant resilience to thermal stress, a hypothesis that warrants further investigation in future studies.

In summary, our study provides an unprecedented, in-depth analysis of the role of DNA methylation in transcriptional reprogramming under abiotic stress. By combining multiple stress conditions—heat, phosphate deficiency, and their combination—across distinct tissues and under experimental setups that closely mimic natural field environments, we offer a comprehensive view of stress-induced epigenetic dynamics. Our findings reveal that transposable element methylation changes do not strongly influence global gene expression, however, changes in intergenic regions may contribute to shaping chromatin accessibility at regulatory regions, potentially modulating transcription factor binding. Moreover, we uncover a specific regulatory effect on a subset of genes within the nuclear mitochondrial DNA region (NUMT), pointing to a possible link between epigenetic remodeling and mitochondrial function. Altogether, these results position DNA methylation as a nuanced layer of regulation that integrates environmental cues into plant adaptive responses. This work sets the stage for future studies aimed at leveraging epigenetic plasticity to enhance crop resilience in the face of increasingly complex stress scenarios.

## Methods

### Plant Material and Growth Conditions

*Arabidopsis thaliana* (Columbia-0 ecotype) was used to study the effects of combined temperature stress and phosphate deficiency. Seeds were surface-sterilized with bleach 50% for 3 minutes, then washed three times and stratified at 4°C for 48 hours before germination. Arabidopsis seedlings were germinated in half-strength Murashige and Skoog (½ MS) medium with vitamins plus 1% sucrose, 1% Difco Agar, and 0.05% MES (pH 5.8). For the low phosphate conditions, phosphate-free MS was used, and KH₂PO₄ was added to reach 30 µM. During the whole experiments, plants were grown in D-Root devices, which allow independent control of root-zone light conditions. During the first 4 days, plants were maintained under control standardized conditions (22°C, 16-hour photoperiod, 60% relative humidity). Each treatment included three biological replicates per tissue type (root and shoot). The TGRooZ system enables precise thermal gradient establishment for root zone studies under controlled growth conditions. This experimental setup maintains root system functionality in plants experiencing elevated aerial temperatures, with preserved physiological performance critical for sustaining shoot biomass production and developmental processes under thermal stress [21].

### Tissue Sampling and Preservation

At the end of the 6 day-treatment period, roots and shoots were separated using sterile scalpels. Tissues were immediately frozen in liquid nitrogen and stored at-80°C until further processing. From the identical material, DNA and RNA were extracted using QIAGEN DNA or RNA kits to ensure high-quality nucleic acids for downstream analyses.

### Whole Genome Bisulfite Sequencing

DNA methylation was analyzed using Whole Genome Bisulfite Sequencing (WGBS). For normal WGBS library constructing, the DNA was fragmented by sonication using a Bioruptor (Diagenode, Belgium) to a mean size of approximately 250 bp, followed by the blunt-ending, dA addition to 3’-end, finally, adaptor ligation (in this case of methylated adaptors to protect from bisulfite conversion), essentially according to the manufacturer’s instructions. Ligated DNA was bisulfite converted using the EZ DNA Methylation-Gold kit (ZYMO). Different Insert size fragments were excised from the same lane of a 2% TAE agarose gel. Products were purified by using QIAquick Gel Extraction kit (Qiagen) and amplified by PCR. At last, Sequencing was performed using the HighSeq4000. Libraries were prepared for paired-end sequencing (150 bp reads) with an average insert size of 300 ± 50 bp. Sequencing was carried out on an Illumina platform with a minimum coverage depth of 30× per sample.

### RNA Sequencing

RNA-seq libraries were prepared from total RNA using standard protocols. Paired-end reads were generated on the Illumina platform. Gene expression reads were aligned to the *Arabidopsis thaliana* TAIR10 genome using Bowtie2 with default parameters, while transposable element (TE) reads were aligned using STAR allowing up to 100 multimapping loci. Read counts for genes and TEs were quantified using HTSeq-count in union mode. Alberto faltaría decir el cut-off (pval y fold Change en cada caso).

## Bioinformatics Analysis DNA Methylation Analysis

### Quality Control and Preprocessing

Initial quality assessment was performed using FastQC v0.12.1. Adapter trimming and quality filtering were conducted with Trim Galore v0.6.10 and Cutadapt v4.3, using the following parameters: minimum quality score of 20 (Phred33), adapter overlap stringency of 1 bp, minimum read length of 20 bp after trimming, and maximum trimming error rate 0.1 (default). Post-trimming quality control was performed to verify adapter removal and overall quality improvement.

### Reference Genome Preparation

The Arabidopsis thaliana TAIR10 reference genome was downloaded from The Arabidopsis Information Resource (TAIR). Bismark genome preparation was performed using bismark_genome_preparation with the HISAT2 aligner option to generate in silico bisulfite-converted genome indices (both C→T and G→A conversions).

### Bisulfite-aware Alignment

Trimmed paired-end reads were aligned to the bisulfite-converted reference genome using Bismark v0.24.2 with HISAT2 v2.2.1 as the underlying aligner and SAMtools v1.3.1 for BAM file handling. Alignment was performed with the --non-directional option to account for non-directional library preparation, --hisat2 to specify the HISAT2 aligner, and --parallel 4 for multi-threaded processing. Alignment efficiency was monitored for each sample, with expected rates >60% for high-quality WGBS data.

### PCR Duplicate Removal

PCR duplicates were identified and removed using deduplicate_bismark in paired-end mode. Deduplication rates ranging around ∼3% were observed across samples, consistent with expected rates for standard WGBS library preparation protocols. Read pairs mapping to identical genomic positions were considered duplicates and removed to prevent artificial inflation of methylation calls.

### Methylation Extraction

DNA methylation data were extracted from deduplicated BAM files using bismark_methylation_extractor with the following parameters: --paired-end for paired-end data processing, --no_overlap to avoid double-counting overlapping read pairs, --comprehensive to extract all cytosine contexts (CG, CHG, CHH), --cytosine_report to generate genome-wide cytosine reports, and --parallel 4 for multi-threading. We used bismark2report afterwards to report the HTML summary of the extraction. Methylation levels were quantified for each cytosine context separately. Quality assessment of methylation extraction confirmed expected methylation patterns for *Arabidopsis thaliana*.

### Differentially Methylated Region (DMR) Identification

DMR analysis was performed using the DMRcaller v1.24.0 package in R/Bioconductor. Biological replicates were pooled by treatment group using readBismarkPool from the methylKit package. Eight experimental groups were defined: Shoot_32_+P (S1-S3), Shoot_32_-P (S4-S6), Shoot_22_+P (S7-S9), Shoot_22_-P (S10-S12), Root_32_+P (S13-S15), Root_32_-P (S16-S18), Root_22_+P (S19-S21), and Root_22_-P (S22-S24).

Pairwise comparisons were conducted to assess temperature effects (32°C TGRooZ vs. 22°C TGRooZ), phosphate deficiency effects (+P vs.-P µM Pi), and effects of combined stresses (32°C TGRooZ-P vs 22°C TGRooZ +P). DMRs were identified using the bins method with the following parameters: 100 bp bin size, score test for statistical significance, p-value threshold of <0.05, minimum of 4 cytosines per bin, minimum methylation proportion difference of ≥0.4, and minimum of 4 reads per cytosine. DMR calling was performed separately for each cytosine context (CG, CHG, CHH) to account for context-specific methylation patterns and regulatory mechanisms in plants.

## RNA-seq Analysis

### Sample Preparation and Sequencing

Total RNA was extracted from *Arabidopsis thaliana* shoot and root tissues corresponding to the same experimental design as described for WGBS analysis: two tissue types (shoot and root), two temperature treatments (32°C TGRooZ and 22°C TGRooZ), and two phosphate conditions (+P and-P), with three biological replicates per condition (24 samples total). RNA-seq libraries were prepared and sequenced on an Illumina platform, generating paired-end 150 bp reads. Raw sequencing data were obtained in FASTQ format.

### Quality Control and Preprocessing

Raw and post trimming sequencing reads were processed using FastQC v0.12.1. to evaluate per-base sequence quality, adapter contamination, sequence duplication levels, GC content distribution, and other quality metrics. Each report is associated with an HTML file for each sample.

### Read Alignment

Gene expression reads were aligned to the Arabidopsis thaliana TAIR10 genome using Bowtie2 with default parameters, while transposable element (TE) reads were aligned using STAR allowing up to 100 multimapping loci. Read counts for genes and TEs were quantified using HTSeq-count in union mode. SAM files were converted to BAM format, coordinate-sorted, and indexed using SAMtools v1.18.

### Gene and TE Expression Quantification

Read counts were assigned to genes using featureCounts from the Subread v2.0.6 package. featureCounts was run with the following parameters:-p for paired-end fragment counting,-t exon to count reads mapping to exonic regions,-g gene_id to aggregate counts at the gene level,-s 2 for reverse-strand-specific counting, and-T 8 for multi-threading. The resulting gene count matrix was used for subsequent differential expression analysis.

Read counts were assigned to genes, exons, and transposable elements (TEs) using htseq-count with the Araport11 annotation GTF file. htseq-count was executed with the following parameters: --mode union to handle overlapping features, --stranded no to specify unstranded counting, --minaqual 0 to include all reads regardless of mapping quality, and --nonunique all to count reads mapping to multiple features. For each BAM file, counts were quantified separately for features of type gene, exon, and transposable_element (using the --type and --idattr options). The resulting count matrices for genes, exons, and TEs were used for downstream differential expression analyses.

### Differential Expression Analysis

Differential gene expression analysis was performed using DESeq2 v1.40.0 in R/Bioconductor. A DESeqDataSet object was constructed from the count matrix and sample metadata, incorporating the factorial experimental design with main effects and interaction terms for tissue type, temperature condition, and/or phosphate condition.

Genes with fewer than 10 reads in at least 3 samples were filtered out to remove lowly expressed genes. DESeq2 normalization and dispersion estimation were performed using default parameters. Reference levels were set to Shoot/Root tissue, 22°C TGRooZ, and +P phosphate for interpretation of results.

Differential expression testing was conducted to identify genes responding to tissue-specific effects, H2 treatment effects, and phosphate deficiency effects. Genes with adjusted p-values <0.05 were considered significantly differentially expressed. Principal component analysis (PCA) was performed on variance-stabilized transformed (VST) data to assess sample clustering and identify potential batch effects.

## Statistical Analysis and Visualization

All statistical analyses were performed in R v4.3.0. Data visualization, including heatmaps, PCA plots, and volcano plots, was generated using ggplot2, pheatmap, and RColorBrewer packages. All analyses were conducted with three biological replicates per condition to ensure statistical power for detecting biologically relevant differences.

Jaccard similarity was analyzed and visualized using https://molbiotools.com. Heatmaps were created using https://heatmapper2.ca

