## Supplementary Figures for "DNA Methylation Dynamics Reveal Unique Plant Responses and Transcriptional Reprogramming to Combined Heat and Phosphate Deficiency Stress"

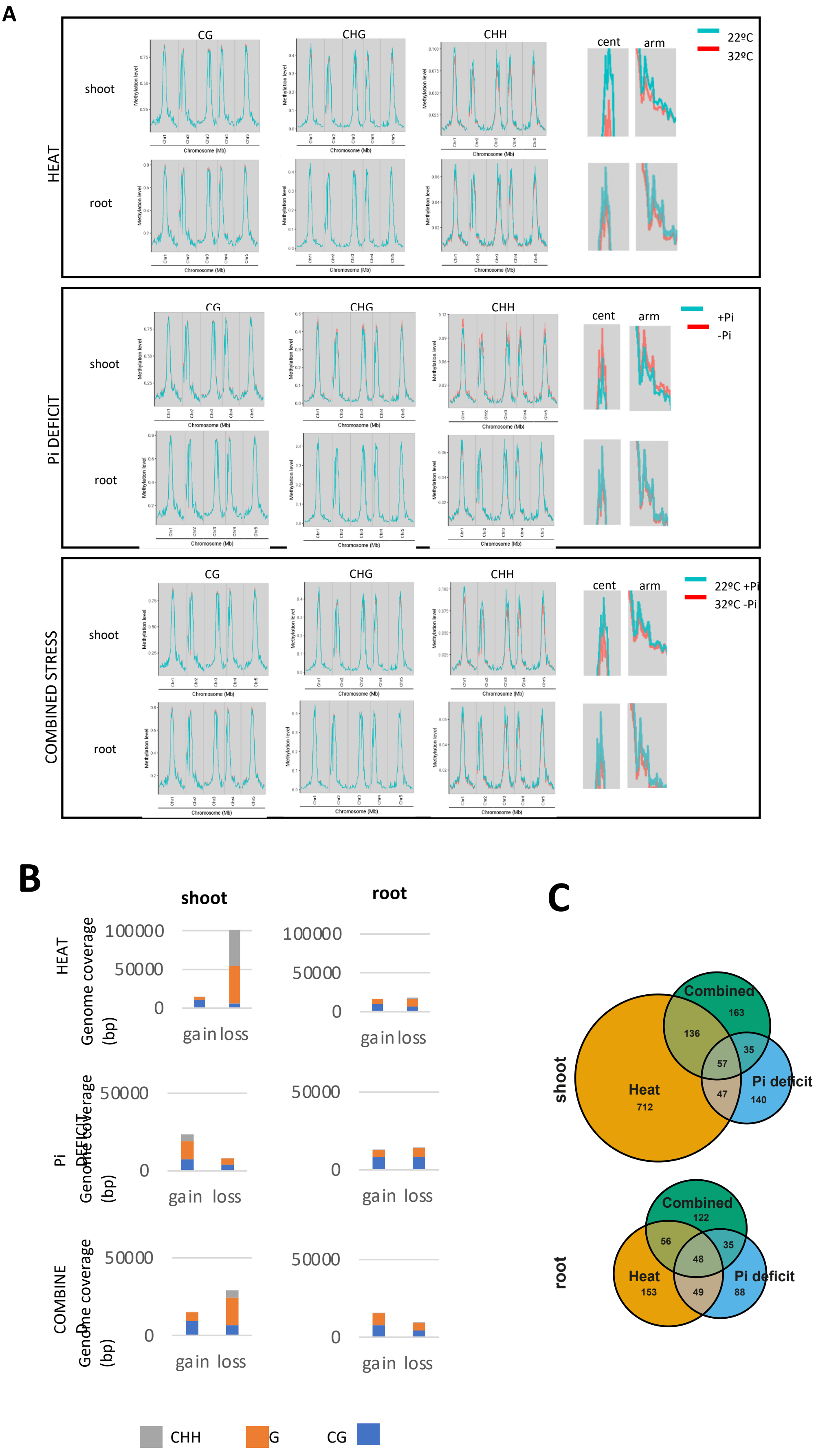

A
HEAT
shoot
root
CG
CHG
CHH
cent
arm
22ºC
32ºC
Methylation level
Chromosome (Mb)
Pi DEFICIT
+Pi
-Pi
COMBINED STRESS
22ºC +Pi
32ºC -Pi
B
shoot
root
HEAT
Genome coverage (bp)
100000
50000
0
gain loss
Pi DEFICIT
COMBINED
CHH
G
CG
C
shoot
Combined
163
136
35
57
Heat
712
47
Pi deficit
140
root
Combined
122
56
35
48
Heat
153
49
Pi deficit
88

**Figure S1. Genome-wide DNA methylation changes under abiotic stresses in Arabidopsis shoots and roots.**

(A) Distribution of methylation levels across CG, CHG, and CHH contexts along the five chromosomes in shoots and roots under heat stress, Pi deficiency, and combined stress compared to control conditions. Insets show a detail of global methylation in centromeres and arms.

(B) Total genomic regions (in base pairs) showing methylation gain or loss in each context (CG, CHG, CHH) for shoots and roots under the three stress conditions.

(C) Venn diagrams illustrating the overlap of elements in differentially methylated regions (DMRs) among stress treatments in shoots (top) and roots (bottom).

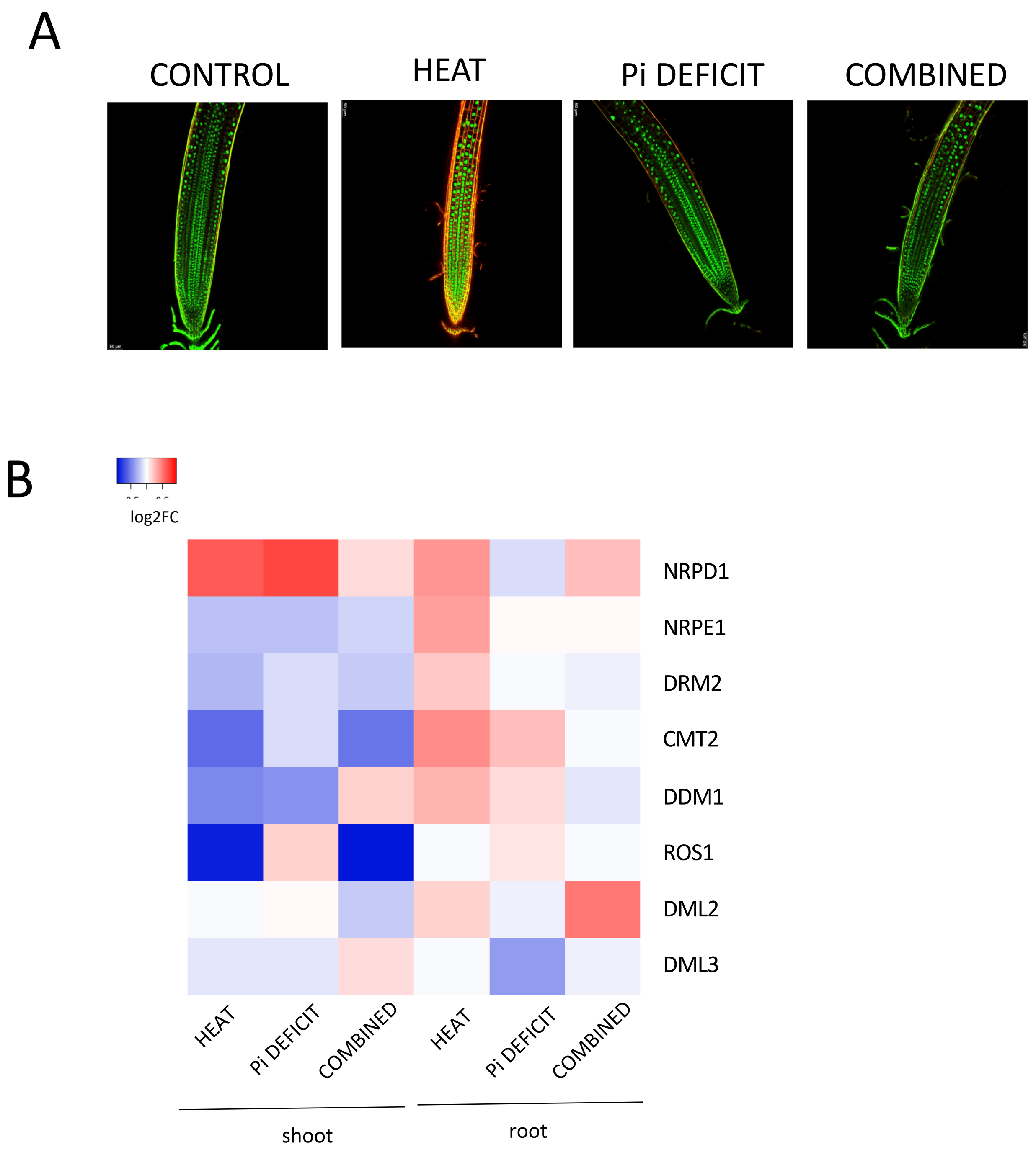
A
CONTROL
HEAT
Pi DEFICIT
COMBINED
B
log2FC
NRPD1
NRPE1
DRM2
CMT2
DDM1
ROS1
DML2
DML3
HEAT
Pi DEFICIT
COMBINED
HEAT
Pi DEFICIT
COMBINED
shoot
root

**Figure S2. Visualization of CMT2-GFP and expression of DNA methylation machinery under abiotic stresses.**
(A) Confocal images of Arabidopsis roots under control conditions, heat stress, Pi deficiency, and combined stress. Fluorescent staining corresponds to CMT2-GFP.
(B) Heatmap showing log2 fold-change in transcript abundance of key genes involved in DNA methylation and demethylation pathways (NRPD1, NRPE1, DRM2, CMT2, DDM1, ROS1, DML2, DML3) in shoots and roots under heat stress, Pi deficiency, and combined stress.

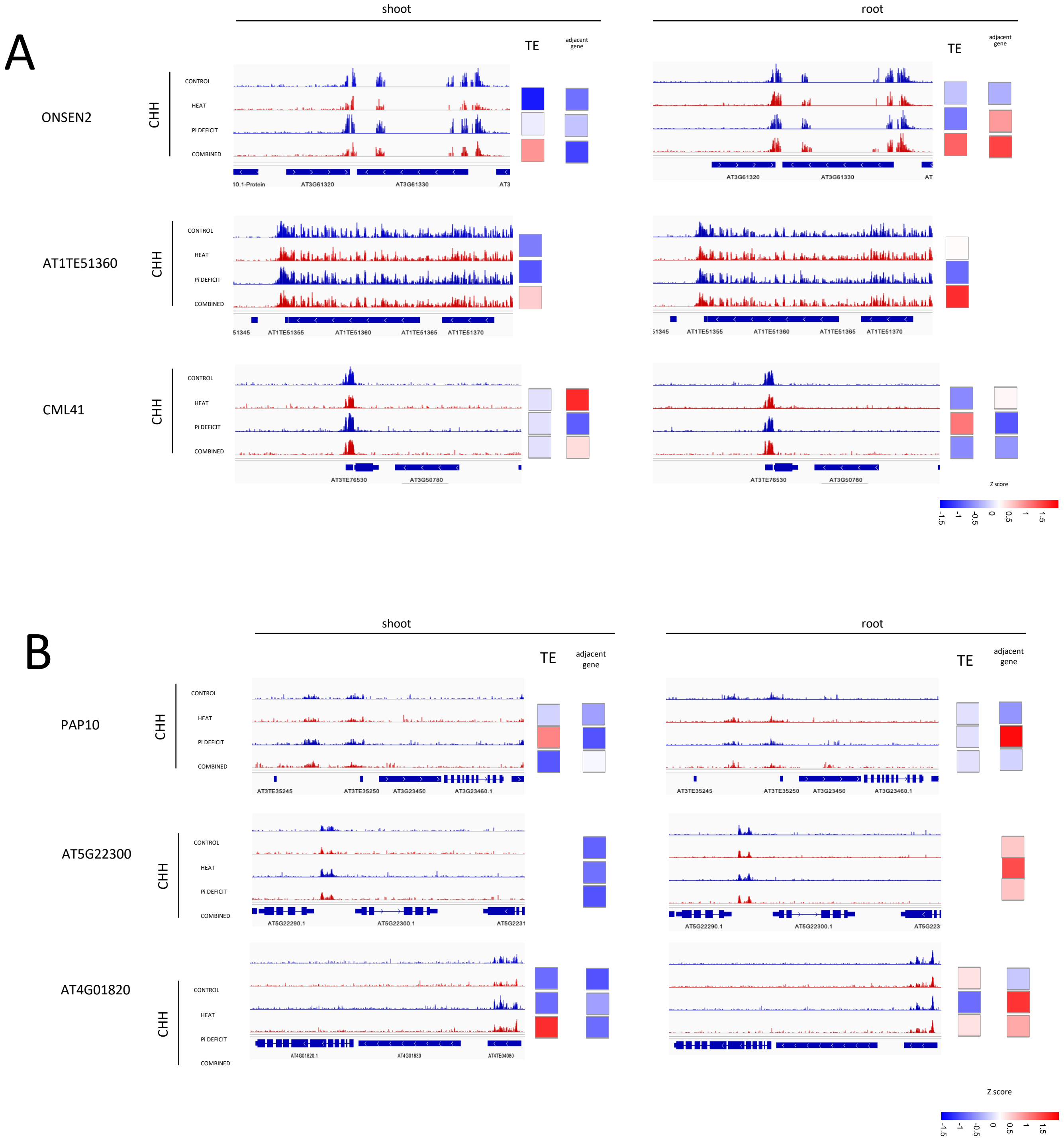
A
shoot
root
TE
adjacent gene
ONSEN2
CHH
CONTROL
HEAT
Pi DEFICIT
COMBINED
AT3G61320
AT3G61330
AT1TE51360
AT1TE51355
AT1TE51365
AT1TE51370
CML41
AT3TE76530
AT3G50780
Z score
-1.5
-1
-0.5
0
0.5
1
1.5
B
PAP10
AT3TE35245
AT3TE35250
AT3G23450
AT3G23460.1
AT5G22300
AT5G22290.1
AT5G22300.1
AT4G01820
AT4G01820.1
AT4G01830
AT4TE04080

**Figure S3. CHH methylation dynamics at transposable elements and adjacent genes under abiotic stresses.**
(A) Genome browser tracks showing CHH methylation profiles for selected loci (ONSEN2, AT1TE51360, and CML41) and expression of TEs and adjacent genes in shoots (left) and roots (right) under control conditions, heat stress, Pi deficiency, and combined stress.

(B) Genome browser tracks showing CHH methylation profiles for selected loci (PAP10, AT5G22300, and AT4G01820) and expression of TEs and adjacent genes in shoots (left) and roots (right) under control conditions, heat stress, Pi deficiency, and combined stress.

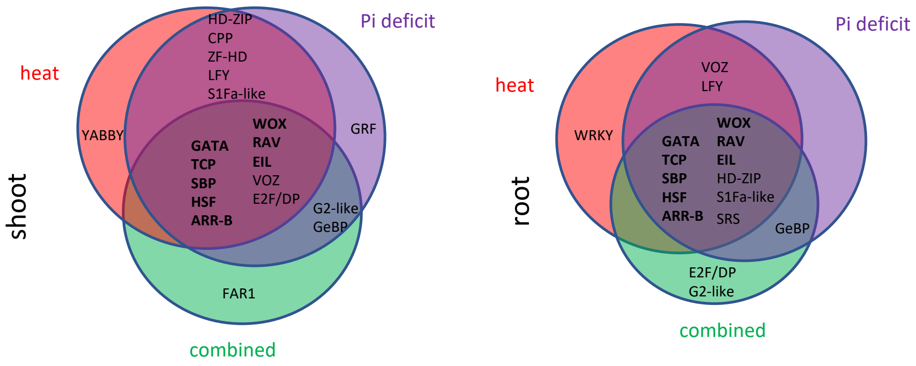
shoot
heat
Pi deficit
combined
HD-ZIP
CPP
ZF-HD
LFY
S1Fa-like
YABBY
GRF
GATA
TCP
SBP
HSF
ARR-B
WOX
RAV
EIL
VOZ
E2F/DP
G2-like
GeBP
FAR1
root
heat
Pi deficit
combined
VOZ
LFY
WRKY
GATA
TCP
SBP
HSF
ARR-B
WOX
RAV
EIL
HD-ZIP
S1Fa-like
SRS
GeBP
E2F/DP
G2-like

**Figure 4. Transcription factor families binding sites in DMRs during stress-specific responses in shoots and roots.**
Venn diagrams showing the overlap of transcription factor binding sites enriched under heat stress (red), Pi deficiency (purple), and combined stress (green) in shoots (left) and roots (right).
